# The hidden state-dependency of transcranial alternating current stimulation (tACS)

**DOI:** 10.1101/2020.12.23.423984

**Authors:** Florian H. Kasten, Christoph S. Herrmann

**Author notes:** Corresponding author: Christoph S. Herrmann, Experimental Psychology Lab, Carl von Ossietzky University, Ammerländer Heerstr. 114 – 118, 26129, Oldenburg, Germany.

## Abstract

Non-invasive techniques to electrically stimulate the brain such as transcranial direct and alternating current stimulation (tDCS/tACS) are increasingly used in human neuroscience and offer potential new avenues to treat brain disorders. However, their often weak and variable effects have also raised concerns in the scientific community. A possible factor influencing the efficacy of these methods is the dependence on brain-states. Here, we utilized Hidden Markov Models (HMM) to decompose concurrent tACS-magnetoencephalography data into transient brain-states with distinct spatial, spectral and connectivity profiles. We found that out of four spontaneous brain-states only one was susceptible to tACS. No or only marginal effects were found in the remaining states. TACS did not influence the time spent in each state. Our results suggest, that tACS effects may be mediated by a hidden, spontaneous state-dependency and provide novel insights to the changes in oscillatory activity underlying aftereffects of tACS.

## 1 Introduction

Non-invasive brain stimulation (NIBS) is increasingly used to assess the involvement of specific brain regions or activity patterns such as neural oscillations for certain cognitive functions. For example, an increasing body of research used interventional approaches to study the role of brain oscillations in cognition by means of rhythmic NIBS such as rhythmic transcranial magnetic stimulation (rTMS) and especially transcranial alternating current stimulation (tACS)^1,2^. In addition, these techniques may offer new avenues for therapeutic interventions^3–5^.

TACS works via the application of weak, alternating currents to the scalp and is believed to engage neural oscillations via entrainment^6–9^. On a physiological level, tACS has repeatedly been shown to alter human brain oscillations for several minutes or even beyond an hour after the offset of stimulation^10–13^. These aftereffects have been suggested to arise due to neural plasticity induced by the stimulation^10,11,14^. In recent years, NIBS techniques received considerable criticism due to their weak and variable effects, leading some authors to question their efficacy altogether^15–18^. It is suspected that a wide variety of individual factors may influence the effectiveness of NIBS^19^, including differences in brain anatomy^20,21^ and genetic dispositions^22^. A couple of studies further suggest, that effects of stimulation may depend on brain-states^13,23–26^. For example, alterations in α-band activity by tACS were observed while participants kept their eyes-open, but were absent when they kept their eyes closed^13,23,25^. These observations were based on experimentally induced, comparably long-lasting changes in the brain’s state. It is, however, likely that, even under relatively constant experimental conditions, the brain spontaneously alternates between different, transient states at rates of a few seconds or even at sub-second ranges, rather than remaining in a constant state^27,28^. Recent work underpins this idea and demonstrated that these transient states exhibit distinct oscillatory activity and connectivity patterns^27,29^. It is thus tempting to assume that these ‘hidden’, spontaneous brain-states may also differ in their susceptibility to the same brain stimulation protocol. In addition, a brain-state perspective may also aid our understanding of the nature of tACS aftereffects. An increase in oscillatory power can in principle occur in multiple ways. It is usually assumed that the effect of tACS results from an increase in the amplitude of the target oscillation (Fig. 1 a,b). However, it is also possible that changes in spectral power arise from a more frequent occurrence of brain-states with comparably higher oscillatory power, without necessarily changing the actual amplitude of oscillations within each state^30^ (Fig. 1c). Alternatively, stimulation effects may be obscured if an increase in power in a given state is accompanied by a reduced occurrence of the state or vice versa (Fig. 1e). In the current study, we aimed to address these questions by adapting a recently proposed analysis pipeline utilizing Hidden Markov Models (HMMs) to decompose neural time-series data into distinct, transient brain-states^27–29^ to two recently obtained tACS-magnetoencephalography (MEG) datasets^20^. We hypothesized that occipito-parietal alpha-tACS differentially affects power in the α-band across brain-states but does not alter the relative occurrence of states.

**Figure 1:**
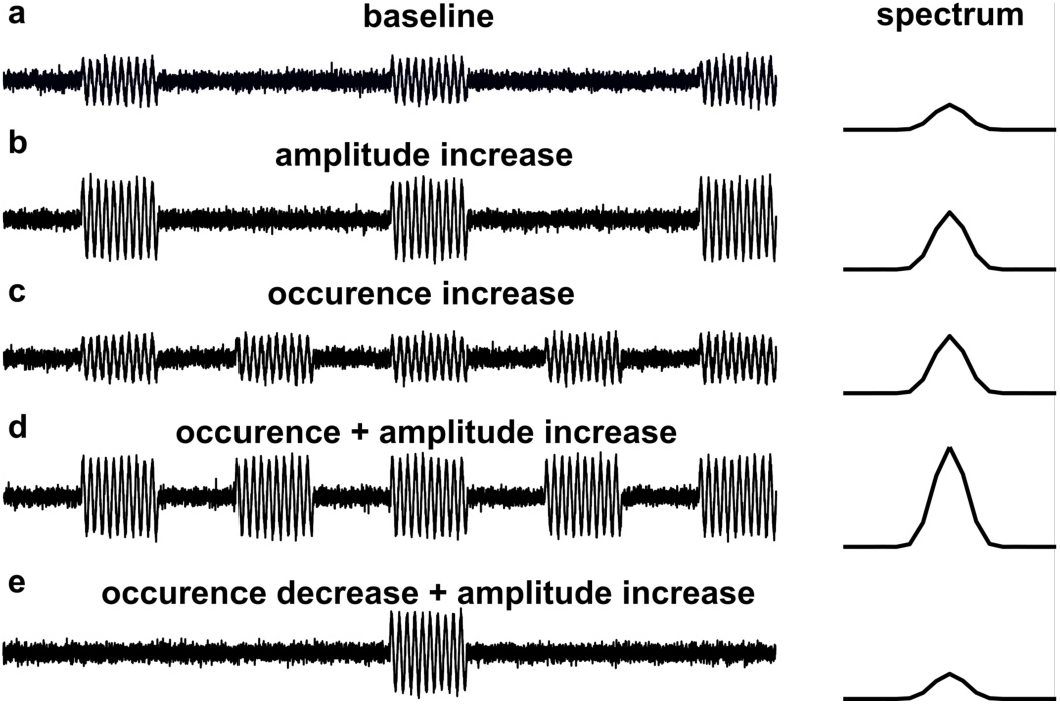
Possible activity changes underlying a tACS effect. **(a)** Hypothetical M/EEG time series before stimulation. The activity pattern alternates between two states: short occurrences of 10 Hz α-activity and ‘no-alpha’ states. The resulting power spectrum is depicted on the right **(b-d)** After stimulation, an increase in the average power spectrum could potentially result from different changes in the activity pattern. **(b)** The amplitude in the alpha-state increases while the occurrence and time-spend in both states remain the same. **(c)** The time spent in the α-state increases, while the time spent in the ‘no-alpha’ state decreases. The amplitude of the oscillation remains the same. **(d)** Both the time-spent in the α-state and the amplitude of the oscillation change. **(e)** The amplitude of the oscillation increases, while the occurrence of the alpha-state decreases. In this scenario, the effect of tACS on oscillatory power might be obscured by the change of state occurrence.

## 2 Results

### 2.1 HMM decomposes time-series into transient states

We reanalyzed data from two recent tACS-MEG experiments^20^. Participants received 20-min of tACS at individual α-frequency (IAF) or sham stimulation. Neuromagnetic activity was acquired for a period of 10-min immediately before and after stimulation. Recordings during stimulation were discarded due to the massive electromagnetic artifact introduced by the stimulation^31^. Signals were projected into source-space by means of a linearly constrained minimum variance (LCMV) beamformer and parcellated into 42 regions of interest (ROIs) covering the entire cortex. Hidden Markov Models (HMMs) were subsequently trained on the data following an approach recently presented in Vidaurre et al.^27^ and a state-wise frequency analysis was carried out, leveraging the probabilistic state sequence returned by the model^29^ (see Fig. 2 for an overview of the analysis). We used the first experiment, carried out as a between subject design (n_tACS_ = 20, n_sham_ = 20), as an exploratory dataset to setup and test the analysis pipeline. The second experiment, performed as a between subject design (N = 19), was used to test whether the obtained results could be confirmed.

**Figure 2:**
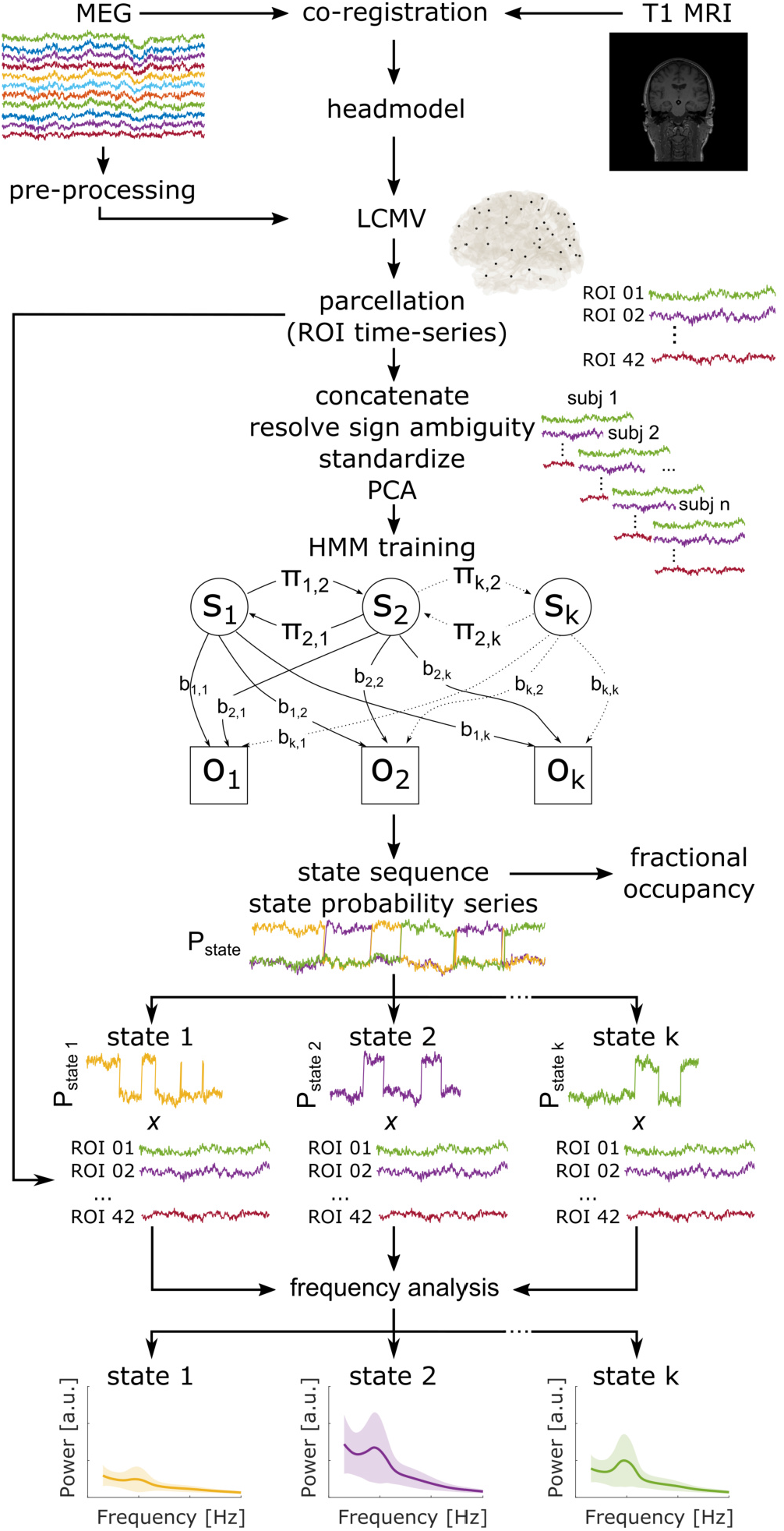
Analysis Pipeline. Preprocessed MEG signals were projected into source-space using an LCMV beamformer. Source time-series were parcellated into 42 ROIs. Before HMM training, single-subject time-series were concatenated, differences in dipole signs across subjects were resolved and the data was standardized within subjects to avoid the HMM to adapt to inter-individual differences. The HMM was then trained on the data in PCA space (84 components). The HMM infers a sequence of a finite number of hidden states (*s_1_, …, s_k_*) based on a set of observable emissions (*o_1_, …, o_k_*). Emissions and states are linked via an emission probability matrix b, where each state has a probability to cause each emission. The transition probability between states is represented in the transition probability matrix π. Both matrices are unknown and need to be estimated from the data by an iterative algorithm (Baum-Welch). The HMM returns the most likely sequence of states, which are used to compute the relative time spent in each state (a.k.a. fractional occupancy), as well as the underlying state probability series. For each timepoint, the series contains a probability of each state being active. By weighting the ROI time series with the state probability series, state-wise frequency spectra and connectivity measures can be computed.

The HMMs decomposed the source time-series into brain-states with distinct spatial, spectral and connectivity profiles. In an initial step, we tested different HMMs with the number of states ranging from two to twelve. After inspection of the results, we decided to settle for a four-state HMM for subsequent analyses, as it returned distinct brain-states without producing redundant (i.e., two or more similar) states (see Supplementary Fig. 1 for an example). The states are in agreement with previous work^27^ and seem to correspond to sensori-motor, visual, and default mode network (DMN) as well a state reflecting posterior suppression (relative to the other states; Fig. 3).

**Figure 3:**
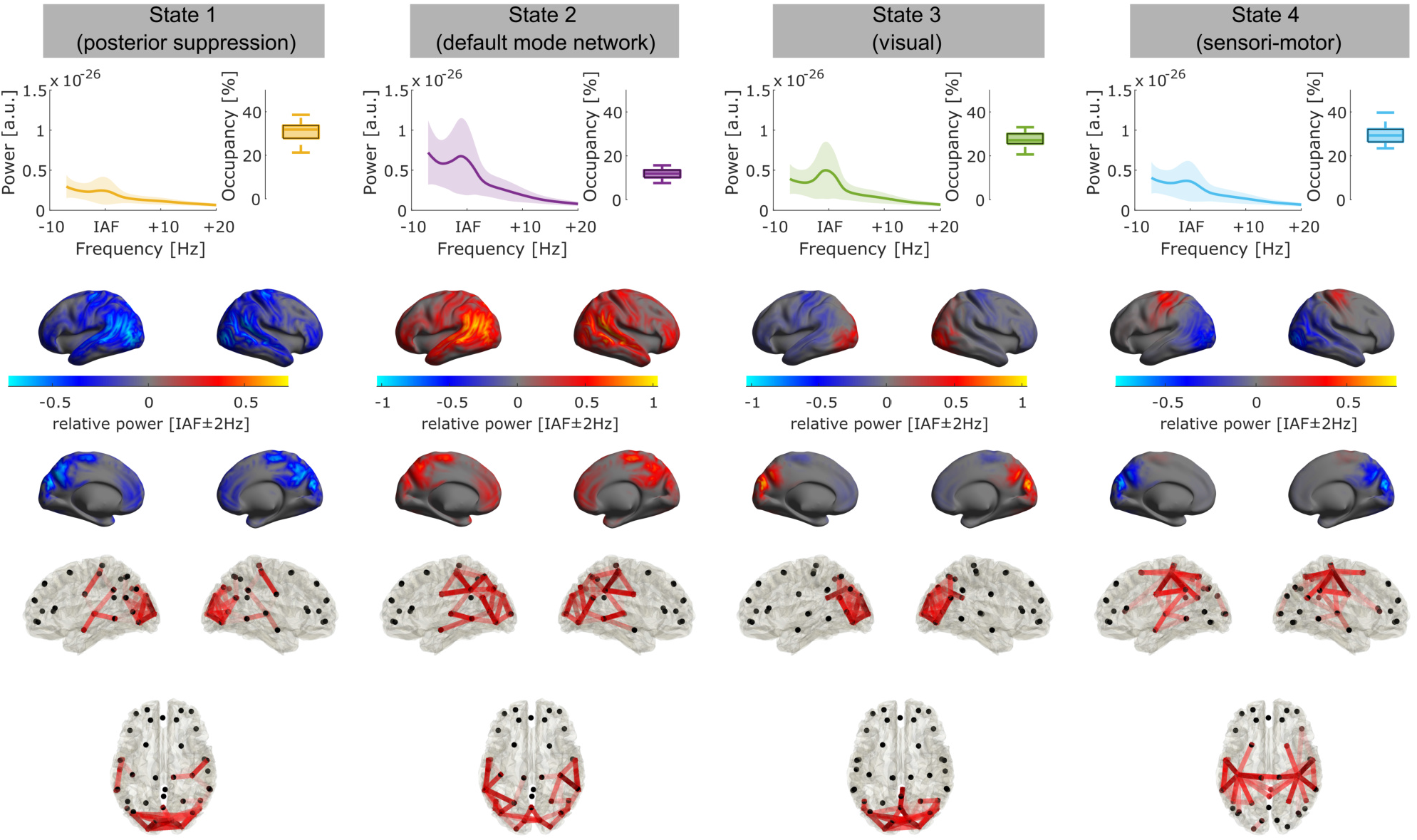
State profiles in experiment 1. Each column depicts power spectra, relative state occupancy (**top row**), spatial maps of α-power relative to the average over all states (**middle row**) and connectivity profiles (**bottom row**, coherence in the α-band) for each of the four states. Connectivity profiles are thresholded to show the 5% strongest connections, darker shades of red indicate higher connection strength. Shaded areas depict standard deviation (S.D.).

### 2.2 tACS differentially affects α-power across states, but not state occupancy

To test if tACS differentially affected brain-states, we compared the power increase in the individual α-band (IAF ± 2 Hz) relative to baseline, averaged over all 42 ROIs, between the tACS and the sham group using a repeated measures ANOVA with the within-subject factor STATE (4-levels) and the between-subject factor STIMULATION (2-levels; tACS vs. sham). Results yielded significant main effects of STIMULATION (F_1,38_ = 9.61, p = .0036, η^2^ = 0.13) and STATE (F_3,114_ = 6.28, p = .008, η^2^ = 0.06) and a significant STIMULATION*STATE interaction (F_3,114_ = 4.28, p = .031, η^2^ = 0.04), indicating that the tACS effect differed across spontaneous brain-states (Fig. 4a,b). To assess which states were affected by stimulation, we subsequently performed non-parametric random permutation cluster t-tests for independent samples comparing the power increase during stimulation and sham in each state. Tests yielded a significant cluster in state 2 (p_cluster_ = .0012, df = 38, Fig. 4c). No clusters were identified in the other states (all p_cluster_ > .14, all df = 38, Supplementary Table 1). We did not find evidence for effects of tACS in the neighboring θ- (IAF-7 – IAF-3) and β-bands (IAF+4 – IAF+20; Supplementary Table 2).

**Figure 4:**
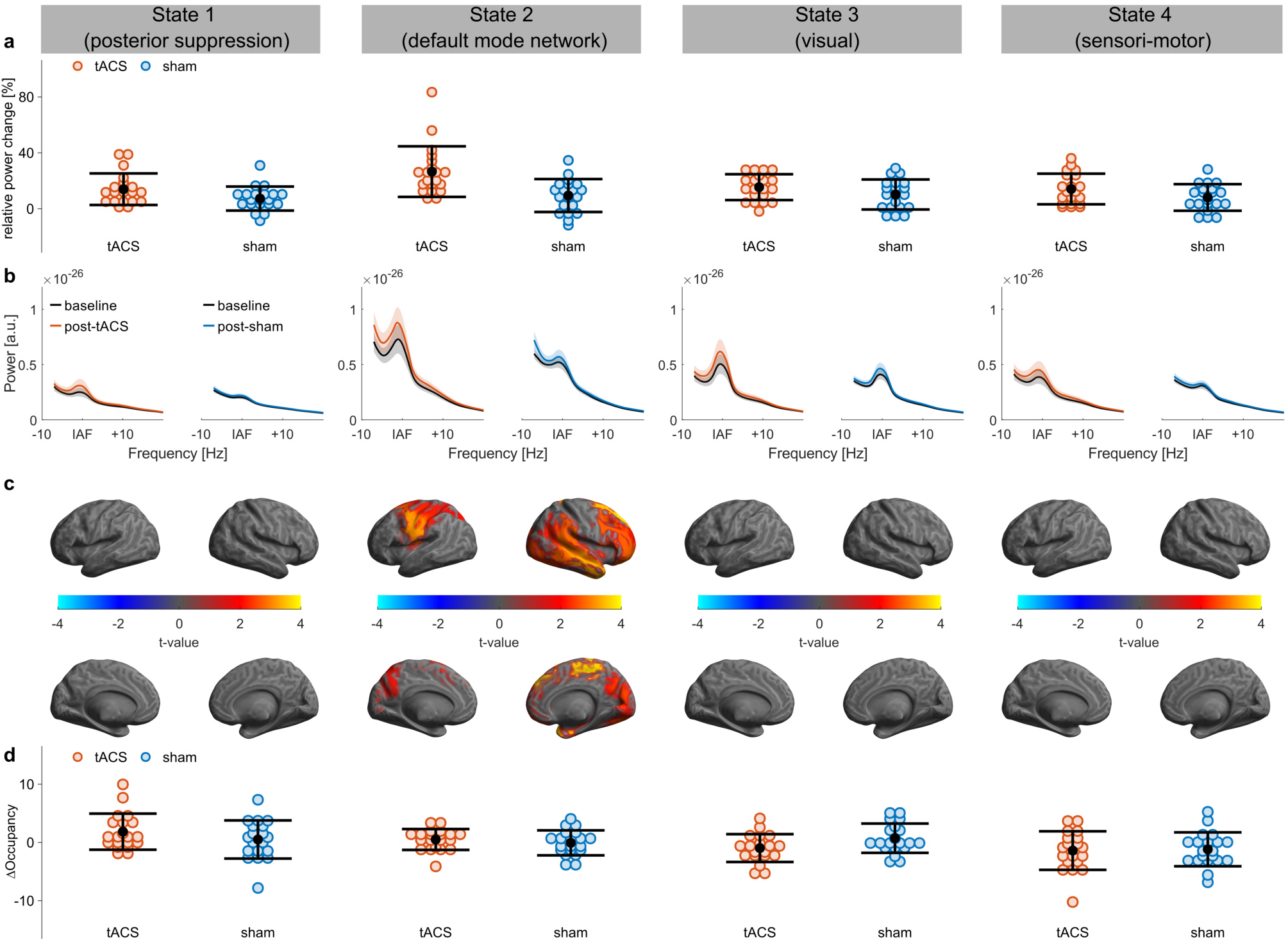
State-specific tACS effect in experiment 1. **(a)** Relative power change per state in the two experimental groups (tACS vs. sham). Black dots and error bars indicate mean and S.D. Colored dots indicate the distribution of individual datapoints. **(b)** Average power spectra over all ROIs before and after tACS or sham stimulation, respectively. Shaded areas depict standard error of the mean (S.E.M.) **(c)** Significant clusters exhibiting larger relative power increase in the α-band after tACS as compared to sham stimulation. T-value maps are thresholded at an α-level of 0.05 (after an additional Bonferroni correction for four multiple comparisons). Only one of the four states (state 2) appeared to be susceptible to tACS. **(d)** Change in fractional occupancy per state for the two experimental sessions (tACS vs. sham). TACS did not change the relative occurrence of states compared to sham.

To test if tACS differentially affected brain-states, we compared the power increase in the individual α-band (IAF ± 2 Hz) relative to baseline, averaged over all 42 ROIs, between the tACS and the sham group using a repeated measures ANOVA with the within-subject factor STATE (4-levels) and the between-subject factor STIMULATION (2-levels; tACS vs. sham). Results yielded significant main effects of STIMULATION (F_1,38_ = 9.61, p = .0036, η^2^ = 0.13) and STATE (F_3,114_ = 6.28, p = .008, η^2^ = 0.06) and a significant STIMULATION*STATE interaction (F_3,114_ = 4.28, p = .031, η^2^ = 0.04), indicating that the tACS effect differed across spontaneous brain-states (Fig. 4a,b). To assess which states were affected by stimulation, we subsequently performed non-parametric random permutation cluster t-tests for independent samples comparing the power increase during stimulation and sham in each state. Tests yielded a significant cluster in state 2 (p_cluster_ = .0012, df = 38, Fig. 4c). No clusters were identified in the other states (all p_cluster_ > .14, all df = 38, Supplementary Table 1). We did not find evidence for effects of tACS in the neighboring θ- (IAF-7 – IAF-3) and β-bands (IAF+4 – IAF+20; Supplementary Table 2).

To test whether tACS affects the time spent in each state (a.k.a. the fractional occupancy), we compared the absolute change in fractional occupancy from baseline to post-stimulation between tACS and sham using a repeated measures ANOVA with the within-subject factor STATE (4-levels) and the between-subject factor STIMULATION (2-levels; tACS vs. sham). The analysis yielded a significant main effect of STATE (F_3,114_ = 4.17, p = .024, η^2^ = 0.09), indicative of a general change of fractional occupancy across states, but neither an effect of STIMULATION (F_1,38_ < 0.01, p = 1, η^2^ < 0.01), nor a STIMULATION*STATE interaction (F_3,114_ = 1.70, p = .25, η^2^ = 0.04), suggesting that tACS did not influence the occurrence of transient states (Fig. 4d).

### 2.3 Validation data

The HMM analysis on the second dataset returned states with spatio-spectral profiles similar to the ones obtained in the first dataset (Fig. 5). Similar to results in the first dataset, the states seem to reflect visual, sensori-motor and DMN activity.

**Figure 5:**
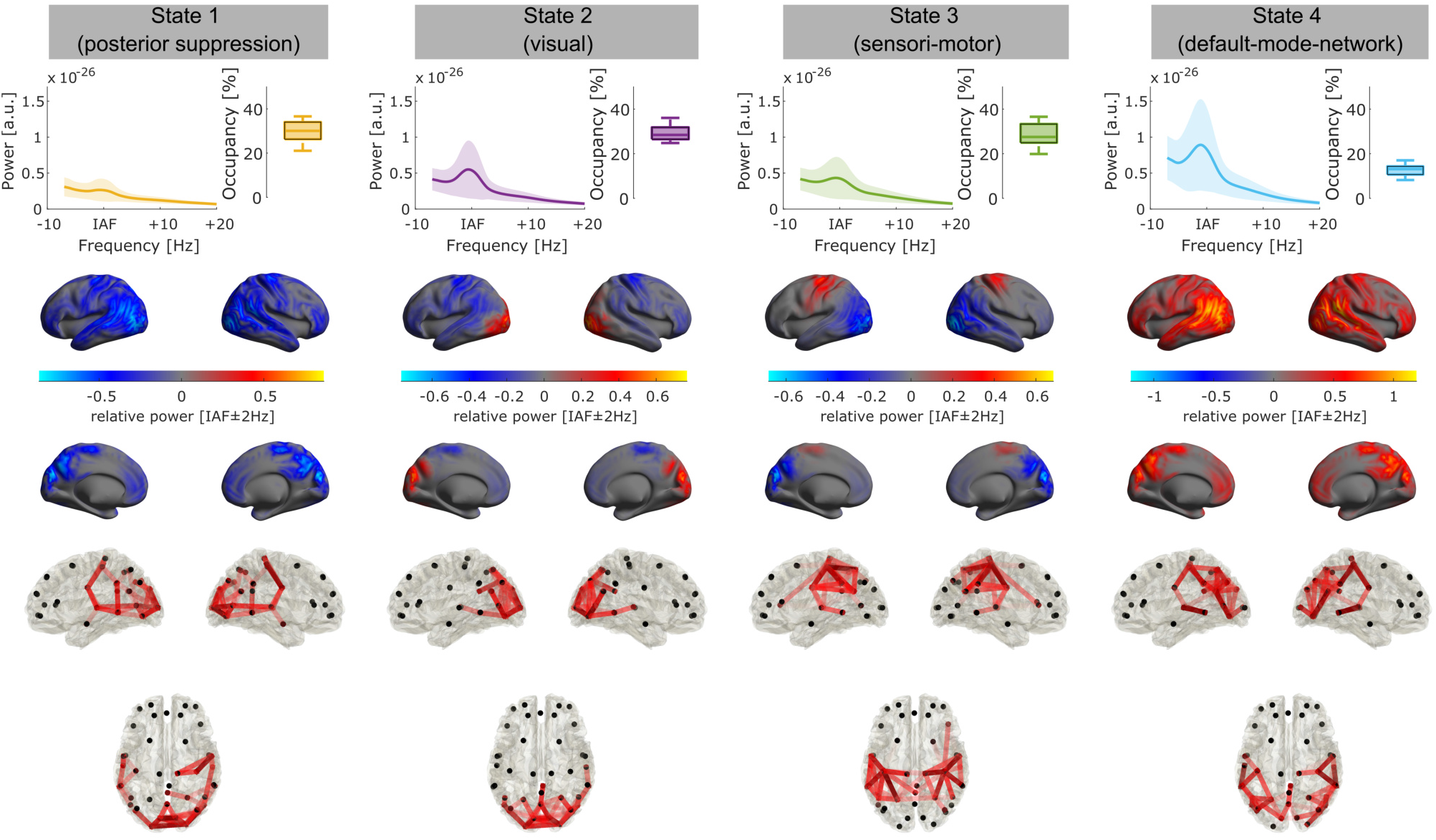
State profiles in experiment 2. Each column depicts power spectra, relative state occupancy (**top row**), spatial maps of α-power relative to the average over all states (**middle row**) and connectivity profiles (**bottom row**, coherence in the α-band) for each of the four states. Connectivity profiles are thresholded to show the 5% strongest connections. Shaded areas depict standard deviation (S.D.)

Again, we assessed the effect of tACS on power in the individual α-band (IAF ± 2 Hz) using a repeated measures ANOVA with the within-subject factors STIMULATION (2-levels; tACS vs. sham) and STATE (4-levels). The analysis yielded significant effects of STIMULATION (F_1,18_ = 9.06, p = .008, η^2^ = 0.06) and STATE (F_3,54_ = 5.05, p = .013, η^2^ = 0.02). Importantly, the STIMULATION*STATE interaction again reached significance (F_3,54_ = 6.88, p = .004, η^2^ = 0.02), replicating the differential effect of tACS across states found in the first dataset (Fig. 6a,b). To assess which states were affected by stimulation, we subsequently performed non-parametric random permutation cluster t-tests for dependent samples comparing the power increase during stimulation and sham in each state. Tests yielded a significant cluster in state 4 (p_cluster_ = .004, df = 18, Fig. 6c) as well as a trend in state (p_cluster_ = .07, df = 18, trends not shown in cluster maps). No effects were found in the other states (all p_cluster_ > .46, df = 18, Supplementary Table 3). Noteworthy, state 4 in this dataset shows strong similarities with state 2 from dataset 1 (Fig. 3 second column, Fig. 5 fourth column), which was the only state showing a significant effect to tACS in the previous analysis (Fig. 4). Again, we did not find evidence for effects of tACS in the neighboring θ- (IAF-7 – IAF-3) and β-band (IAF+4 – IAF+20; Supplementary Table 4).

**Figure 6:**
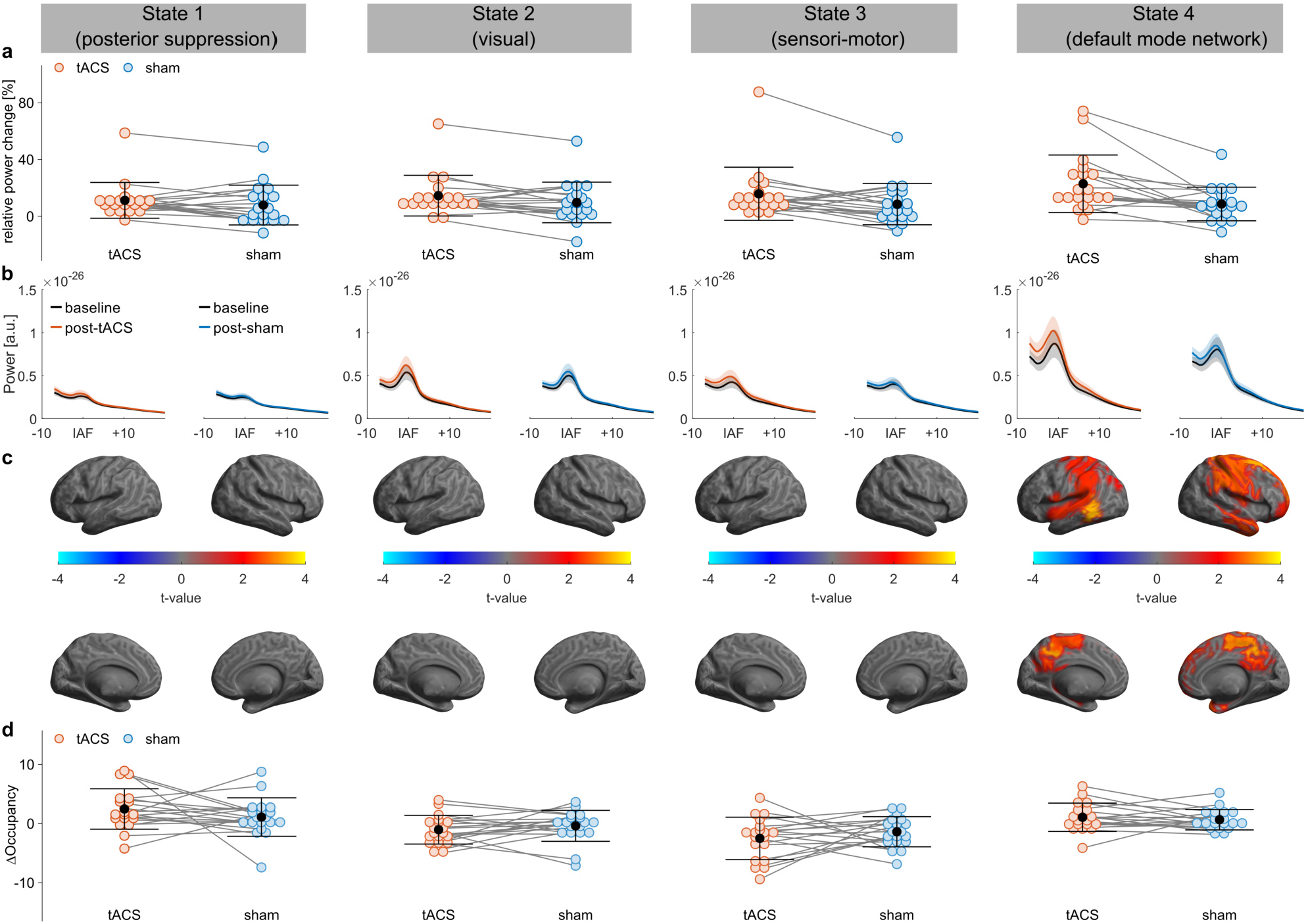
State-specific tACS effect in experiment 2. **(a)** Relative power change per state in the two experimental groups (tACS vs. sham). Black dots and error bars indicate mean and S.D. Colored dots indicate the distribution of individual datapoints. Gray lines indicate within subject differences between stimulation sessions **(b)** Average power spectra over all ROIs before and after tACS or sham stimulation, respectively. Shaded areas depict standard error of the mean (S.E.M.) **(c)** Significant clusters exhibiting larger relative power increase in the α-band after tACS as compared to sham stimulation. T-value maps are thresholded at an α-level of 0.05 (after Bonferroni correction for four multiple comparisons). Only one of the four states (state 4) appeared to be susceptible to tACS. **(d)** Change in fractional occupancy per state for the two experimental sessions (tACS vs. sham). TACS did not change the relative occurrence of states compared to sham.

We next assessed whether tACS affected the time spent in each state by submitting the absolute change in fractional occupancy from baseline to the post-stimulation period to a repeated measures ANOVA with the within subject factors STIMULATION (2-levels, tACS vs. sham) and STATE (4-levels). Once again, the analysis revealed a significant effect of STATE (F_3,54_ = 10.25, p < .001, η^2^ = 0.21), but neither a main effect of STIMULATION (F_1,18_ < 0.01, p = 1, η^2^ < 0.01), nor a STIMULATION*STATE interaction (F_3,54_ = 1.07, p = .33, η^2^ = 0.03), suggesting that tACS did not affect the relative time spent in each of the states (Fig. 6d).

## 3 Discussion

In the current study, we aimed to reveal the influence of hidden, spontaneous brain-state dynamics on the effects of tACS. This end, we adapted a novel analysis framework to decompose neural time series data into brain-states using Hidden Markov Models^27–29^. These states are characterized by distinct spatial, spectral and connectivity profiles. The approach further allowed us to gain insights to the nature of changes underlying the increase in spectral power often associated with the tACS^11,12,32^. Our results suggest that tACS distinctly and consistently modulated α-power in the same state across both datasets, which we identified as most likely reflecting DMN activity. Other states were not or only marginally affected. We did not find evidence that the time spend in any of the states was modulated by tACS, or that the stimulation gave rise to entirely new brain-states.

TACS is generally considered a subthreshold stimulation approach. As such, it is believed to be capable of modulating pre-existing oscillatory activity, but not of inducing new oscillatory activity. In line with this idea, the state most affected by stimulation was one that shows considerable amounts of endogenous activity in the alpha-band. To our surprise, however, the state that captured visual alpha-oscillations, seemed to have not been affected by tACS. The Cz-Oz montage used during the experiment was originally designed to maximize current in the occipital lobe^33^. This raises the question, which properties of a brain-state determine, whether it is susceptible to tACS. For example, it has been suggested that states with already high oscillatory activity cannot be further elevated due to ceiling effects^13,23^. Alternatively, it has been argued that at least some involvement of the target oscillation in the state is necessary for the stimulation to elicit effects^24,34^. Unfortunately, neither of these explanations can fully account for our findings of a susceptible, high alpha-power DMN state and an unsusceptible high α-power visual state. This may indicate that our current perspectives on the state-dependency of tACS effects may be too simplistic. To what extent other features of brain-states such as the number of brain regions involved, their connectivity profiles, the duration for which the state is active, the nature of the underlying information processing or subtle differences in the underlying activity patterns might play a role remains largely elusive. Noteworthy, the state most affected by tACS in both datasets happens to be the one that was active for the least amount of time, while exhibiting the most widespread activation and connectivity profile.

The existence of a ‘hidden’ spontaneous state dependency of tACS effects has implications for the variability of stimulation effects. As illustrated in Fig. 1, changes in the time spend in the brain-state susceptible to stimulation may obscure stimulation effects. Although we did not find evidence that tACS induces such changes in our data, it may be possible that they may occur at random or due to factors unrelated to stimulation (e.g., fatigue/time on task, context conditions, experimental manipulations, etc.). Another aspect that may be worth considering, is the effective stimulation duration that may follow from our results. The susceptible brain-state is active for <20% of the recording time (even if we take a liberal stance and include the state with a trend in experiment 2, we would still be well below 50%). Assuming that the state dynamics during tACS remain comparable to those before and after stimulation, and that tACS only exerts its effects during the time the susceptible state is active, this would imply that our 20-min period of tACS may have contained less than 4-min of effective stimulation. It should be noted, though, that these considerations are highly speculative as the HMM approach cannot be applied during stimulation due to the contamination of MEG signals with a large electromagnetic artifact^31^.

As with all scientific research, some limitations of the current study and its methodology deserve discussion. The HMM requires the user to pre-specify the number of states to-be found in the data. In our analysis we settled for a four state HMM, as larger numbers resulted in a certain redundancy (i.e., similarity between two or more states) of the states returned by the model. This does, however, not imply that the brain exclusively switches between these four states. Specifying a larger number of states, may be useful depending on the specific research question, e.g., if a more fine-grained decomposition of states is desired. For example, using a 12-state HMM could be used to identify subnetworks of the DMN^27^. However, such more detailed descriptions of the data come at the cost of a larger number of multiple comparisons that have to be taken into account and may hamper statistical power and an overall more complex pattern of results that may be more difficult to interpret. The results at hand were obtained from a re-analysis of two existing datasets^20^. Consequently, they cannot be interpreted as an independent replication of the aftereffect of tACS. Further, the datasets only provide data for one stimulation montage and frequency along with a sham control. Future studies adapting the HMM approach to experiments with different stimulation protocols may thus be desirable to better generalize the current findings. One could, for example, conceive stimulation montages to specifically target each of the states reported here, to differentially test their susceptibility to stimulation. For the current study, we refrained from such more complex designs, as we aimed to first establish that transient brain-states identified by HMMs differ in their response to tACS at all. Another interesting application of HMMs in the context of tACS effects may additionally lie in the combination with cognitive tasks. Given a sufficiently complex number of states, HMMs may allow to uncover which subprocess of a task is affected by stimulation. This may foster our understanding of how tACS exerts its behavioral effect, or to even tailor stimulation more specifically to target such subprocesses.

In general, HMMs may have a wide range of applications in the research of non-invasive brain stimulation. Their usage is not limited to M/EEG signals, but they can be utilized in any neuroimaging modality. For example its application to fMRI^28^ might be particularly interesting to study spontaneous state dependency of tDCS. HMMs thus offer a versatile tool to uncover hidden dynamics behind brain stimulation effects and could foster our understanding of brain-state-dependency by providing a more detailed view on neuroimaging data of brain stimulation experiments.

## 4 Methods

### 4.1 Participants

All analyses in the current study were carried out on two pre-existing tACS-MEG datasets^20^. The first one was carried out as a between subject design on 40 participants (24 ± 3 years, 20 females) with participants randomly assigned to one out of two experimental groups (tACS vs. sham), while counterbalancing for participants’ sex. The second experiment was performed as a within-subject design (N = 19, 25 ± 3 years, 11 females), with participants receiving both stimulation conditions at random order across two experimental sessions scheduled on two separate days. Participants were right-handed according to the Edinburgh Handedness-Inventory^35^, had normal or corrected-to-normal vision, were non-smokers, without history of neurological or psychiatric disease and medication-free at the day of the experiment. Written informed consent was obtained from all subjects, both experiments were approved by the Commission for Research Impact assessment and Ethics at the University of Oldenburg.

### 4.2 Magnetoencephalogram (MEG)

MEG signals were obtained at a rate of 1 kHz using a 306-channel whole-head MEG system (Elekta Neuromag Triux System, Elekta Oy, Helsinki, Finland), inside a magnetically shielded room (MSR; Vacuumschmelze, Hanau, Germany). Participants head-position was continuously monitored using five head-position indicator (HPI) coils, attached to participants’ heads. Positions of the coils were digitized along with participants’ head shapes with a Polhemus Fastrak (Polhemus, Colchester, VT, USA) for later co-registration with structural MRIs (T1-weighted 3D sequence, MPRANGE, TR = 2000 ms, TE = 2.07 ms, slice thickness: 0.75 mm; Siemens Magnetom Prisma 3 T MRI machine, Siemens, Erlangen, Germany).

### 4.3 Electrical Stimulation

TACS was administered via two surface-conductive rubber electrodes positioned over locations Cz (7 × 5 cm) and Oz (4 × 4 cm) of the international 10-10 system via an electrically conductive, adhesive paste (ten20 paste, Weaver & Co, Aurora, CO, USA). The stimulation waveform was generated using a constant current stimulator (DC Stimulator Plus, Neuroconn, Illmenau, Germany), remote-controlled via a digital-to-analog converter (NI-USB 6251, National Instruments, Austin, TX, USA) and a Matlab script (MATLAB 2016a, The Math Works Inc., Natick, MA, USA). Stimulation currents were guided into the MSR via the MRI extension-kit of the stimulator. Impedances were kept below 20 kΩ, including two 5 kΩ resistors inside the stimulation cables. Participants received 20 min of active tACS or sham stimulation (30-sec of tACS at the beginning of the stimulation period) at their individual α-frequency IAF. IAF was determined from a 3 min resting MEG obtained prior to the main experiment. 10-min of MEG were recorded directly before and after stimulation. To ensure participants remained attentive and kept their eyes open during the recording, they were asked to perform a simple visual change detection task. To this end, a white fixation-cross on gray background was rear-projected onto a screen (distance: ~100 cm) inside the MSR. Participants had to manually respond to a 500 ms rotation of the cross by 45°, occurring at random intervals with an SOA of 10-110-sec. Additional details on the experimental procedures, including debriefing and the assessment of adverse effects can be found in a previous publication based on the data^20^

### 4.4 Data analysis

#### 4.4.1 Preprocessing

MEG preprocessing was performed in MATLAB 2019b using the fieldtrip toolbox^36^. HMM training and state analysis was performed using the HMM-MAR toolbox (https://github.com/OHBA-analy-sis/HMM-MAR). Statistical analysis was carried out in R 3.6.1 (The R Core Team, Vienna, Austria) running on R-Studio (RStudio Team, PBC, Boston, MA, USA). Statistical functions provided by the fieldtrip toolbox were utilized to compute cluster-permutation statistics.

In a first step, spatio-temporal signal-space-seperation (tSSS) was applied to the data to suppress external interferences and correct for head-movements during the recording^37–39^. TSSS was applied using MaxFilter™ v2.2 (Elekta Neuromag Oy, Helsinki, Finland) with standard settings (Lin = 8, Lout = 3, correlation limit = .98). Signals were subsequently imported to Matlab. To reduce computational demands, the analysis was exclusively performed on the 102 magnetometer channels. Signals were resampled to 256 Hz and filtered between 1 Hz and 40 Hz using 6^th^-order forward-backward Butterworth filters. An independent component analysis (ICA) was performed to remove signal components reflecting eye-movements, heartbeat and stimulator noise. MEG time-series were then projected into source-space using an LCMV beamformer^40^ with a 10 mm source-grid warped into Montreal Neurological Institute (MNI) space. Source time series were subsequently parcellated into 42 virtual channels representing the activity of cortical regions of interest (ROIs) covering the entire cortex. The regions correspond to the ROIs used in ref^27^. A weight-matrix was used to project data into brain-space for visualization purposes. A multivariate correction for spatial-leakage^41^ was applied using the ROInets toolbox (https://github.com/OHBA-analysis/MEG-ROI-nets). The correction reduces artificial correlations between adjacent virtual channels, while resembling the original data as close as possible. Such correlations are known to bias connectivity estimates and can hamper HMM training. A conceptual overview of the analysis pipeline is provided in Fig. 2.

#### 4.4.2 Hidden Markov Model

HMMs are used to infer a sequence of a finite number of hidden states of a system, based on their observable emissions (e.g., patterns of measurable brain signals), where each state has a certain probability to be accompanied by each emission. It is assumed that at each timepoint *t,* the system is in one out of *K* discrete states and that the observable data *o* at *each* timepoint are drawn from an observation distribution, which is of the same family in all states. However, for each state the observation model has a different set of parameters (e.g., different mean and/or SD for a Gaussian observation model)^27^. The transition probability between states is Markovian, i.e., it depends exclusively on the previous state *s_t−1_*:

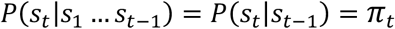

where *π_t_* is the (*K × K*) transition probability matrix, containing the transition probability between all states. The standard approach for training an HMM with a fixed number of states and unknown transition and emission probabilities is the Baum-Welch algorithm (also referred to as the Forward-Backward algorithm)^42^. The algorithm estimates both the emission and transition probabilities by starting off with an initial estimate and then iteratively optimizing both probability matrices. To facilitate the handling of our relatively large dataset, we opted to run the algorithm with a recently developed stochastic inference procedure implemented in the HMM-MAR toolbox that allows to perform the optimization using computationally more efficient subsets of data^43^. Once the transition and emission probabilities are estimated, a state probability sequence *γ*, indicating the probability of each state being active at each timepoint, can be computed along with the most likely state sequence (Fig. 2).

We followed the analysis framework described by Vidaurre et al.^27^ using a time-delay-embedded HMM (TDE-HMM) trained on the concatenated ROI time series of all participants. This variant of the HMM, uses a Gaussian observation model with zero mean to model the data over a certain time window, effectively utilizing the data autocovariance across regions. The TDE-HMM is sensitive to changes in power and phase-locking while being able to deal with a relatively large number of channels^27^. We chose a TDE-HMM with an observation window of 15 samples centered around each *t* and working in PCA-space with 84 components (explaining ~68% of the variance in both datasets). Recordings from all participants were concatenated before training the HMM. This forces the model to return a common set of states across the whole dataset. To avoid that the model assigns states based on inter-individual differences between subjects, ROI time-series were standardized within each subject and the dipole-sign-ambiguity of individual time-series was resolved using the approach presented in ref^27^. We visually confirmed that each state was present in each subject (Supplementary Fig. 3,4).

A common challenge in the application of HMMs, particularly to biological data, is the definition of the number of states for the model, which have to be pre-specified by the user. Objective model selection procedures, e.g., based on Akaike’s or Bayesian Information Criterion (AIC, BIC), or the comparison of free energy^28^, have been proposed for the selection of model complexity. However, these approaches often favor models with large numbers of states that are difficult to handle and interpret^44^. It is thus suggested to take a more practical approach integrating objective measures with practical considerations such as interpretability of results and usefulness in the light of the research question^44^. Indeed, the free energy for our eleven HMMs monotonically decreased from the 2- to the 12-state HMM (Supplementary Fig. 2). However, two “knees” in the trajectory indicate that after including state 4, additional states have less impact on free energy as compared to the previous states. In addition, HMMs with larger numbers of states tended to return states that are similar with respect to their spatial and spectral patterns (e.g., the 6-state HMM, Supplementary Fig. 1), indicating that a state may have been split into sub-states with only minor differences, which is undesirable for our analysis. Based on these considerations we decided to settle for a 4-state HMM for subsequent analyses.

#### 4.4.3 Fractional occupancy and state-wise frequency analysis

After HMM training, we utilized the obtained probabilistic state sequence to compute the fractional occupancy, i.e., the relative time each subject spent in each state, during the baseline and post-stimulation blocks, respectively. To this end, each time-point was assigned to the most probable state and the proportion of time points relative to the total amount of time points per subject and block was computed for each state.

In order to obtain state-wise frequency and connectivity profiles, we leveraged the probabilistic state-time course to perform a state-wise multi-taper analysis (Fig. 2, bottom)^29^. The standard multi-taper power spectral density of the entire time series is given by |*S(f)*|^2^, with

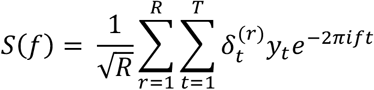

where 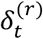 is the value of the *r^th^* taper at time point *t*. To obtain a state-wise spectral analysis for each of the *k* states, the data *y* at each time point *t* is weighted by the probability of being in that state *ρ*, such that the PSD for the *k^th^* state can be obtained by:

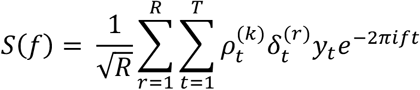

with

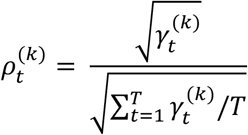

where the normalization term for 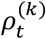 is chosen to preserve the total power of the signal, i.e. the sum of the state spectra corresponds to the total spectrum of the signal^29^. From these PSD estimates, measures of connectivity, such as coherence can be computed.

State-wise PSD and coherence were computed separately for the baseline and post-stimulation blocks using Slepian sequences with 7 tapers and a time-bandwidth product of 4. Importantly, we used the unstandardized ROI time-series for the analysis as the standardization can cancel out differences between blocks and subjects (e.g., stimulation effects). For inspection of the states’ overall spectral, spatial and connectivity profiles, results were averaged across recording blocks and subjects, after re-aligning individual spectra on the IAF. For statistical analysis, PSD in each recording block was averaged within the individual α-band (IAF ± 2 Hz). To facilitate statistical comparisons, the relative change in IAF band power after stimulation relative to baseline was computed:

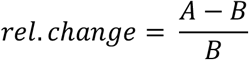

where *A* is IAF band power after tACS or sham stimulation, and *B* is IAF band power in the baseline recording.

#### 4.4.4 Statistical analyses

The power change with respect to baseline, averaged over all 42 ROIs, as well as the change in fractional occupancy were assessed using repeated measures ANOVAs with factors STIMULATION (2-levels, tACS vs. sham, dataset 1: between-subject, dataset 2: within-subject) and STATE (4-levels, within-subject in both datasets). Greenhouse-Geisser corrected p-values are reported if sphericity was violated.

To resolve in which states tACS led to a larger power increase relative to baseline as compared to sham, relative power changes in each state of all 42 ROIs were submitted to non-parametric random permutation cluster t-tests (dataset 1: independent samples, dataset 2: dependent samples), using 10,000 randomizations and Monte-Carlo estimates for p-values. The obtained p-values were Bonferroni-corrected for the 4 multiple comparisons in addition to the cluster correction within each test.

## 5 Acknowledgements

This research was supported by the Neuroimaging Unit of the Carl von Ossietzky University Oldenburg funded by grants of from the German Research Foundation (3T MRI INST 184/152-1 FUGG and MEG INST 184/148-1 FUGG). Christoph S. Herrmann was supported by a grant of the German Research Foundation (Deutsche Forschungsgemeinschaft, DFG) under Germany's Excellence Strategy – EXC 2177/1 - Project ID 390895286.

## 6 Author Contributions

FHK and CSH conceived the study; FHK analyzed the data, FHK and CSH wrote the manuscript.

## Supplementary Information

### Supplementary Figures

**Supplementary Figure 1:**
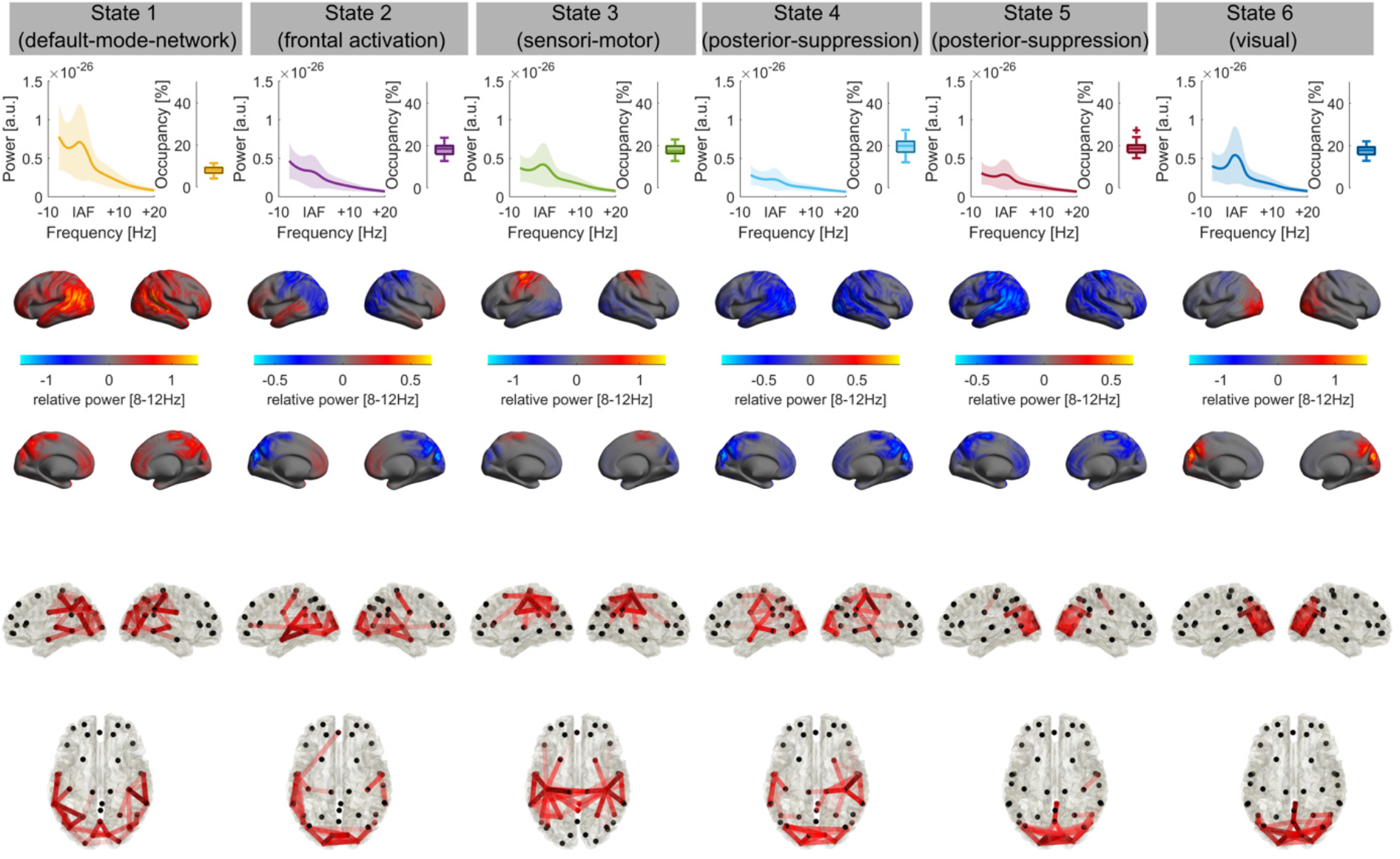
State profiles in experiment 1 for a six state HMM. Each column depicts power spectra, relative state occupancy (**top row**), spatial maps of alpha power relative to the average over all states (**middle row**) and connectivity profiles (**bottom row**, coherence in the alpha band) for each of the six states. Connectivity profiles are thresholded to show the 5% strongest connections, darker shades of red indicate higher connection strength. Shaded areas depict standard deviation (S.D.). The model returned two relatively similar states (4 and 5), showing similar spatial patterns in the alpha band and strong connectivity in occipital regions (although with some central connections in state 4).

**Supplementary Figure 2:**
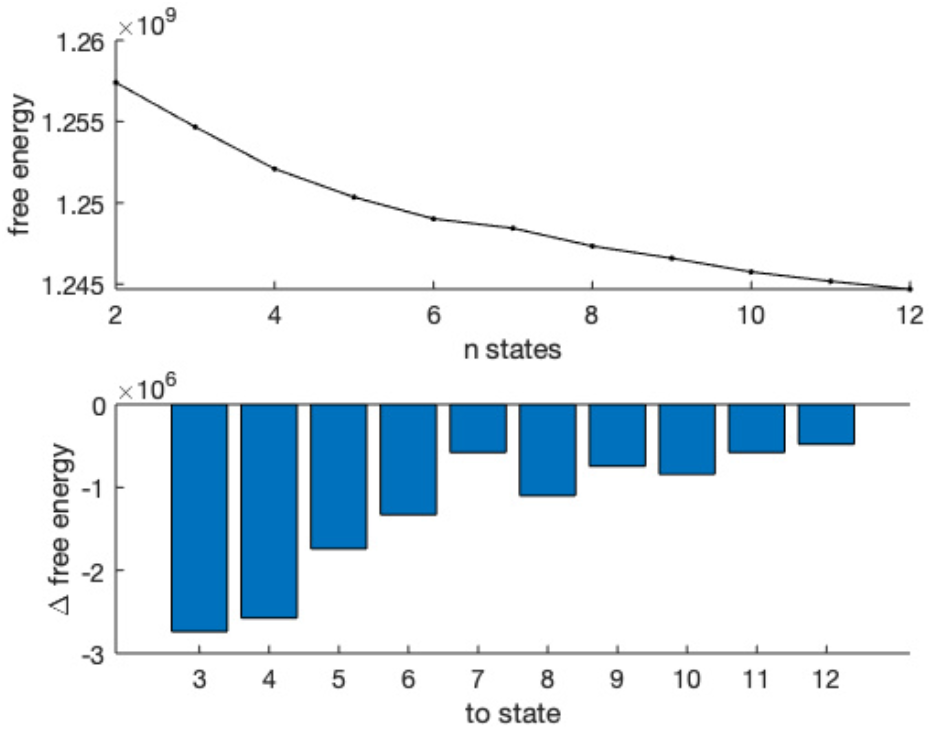
Change in free energy per HMMs as a function of states. Free energy for eleven different HMMs is shown (n_states_ ranges from 2 to 12). Top panel shows the absolute free energy per n_states_, bottom panel shows the change in free energy from one HMM to the next. Free energy monotonically decreases as more states are included in the HMM. However, there are two “knees” in the trajectory, indicating that after the inclusion of state 4 and 6, the benefit of more states starts declining.

**Supplementary Figure 3:**
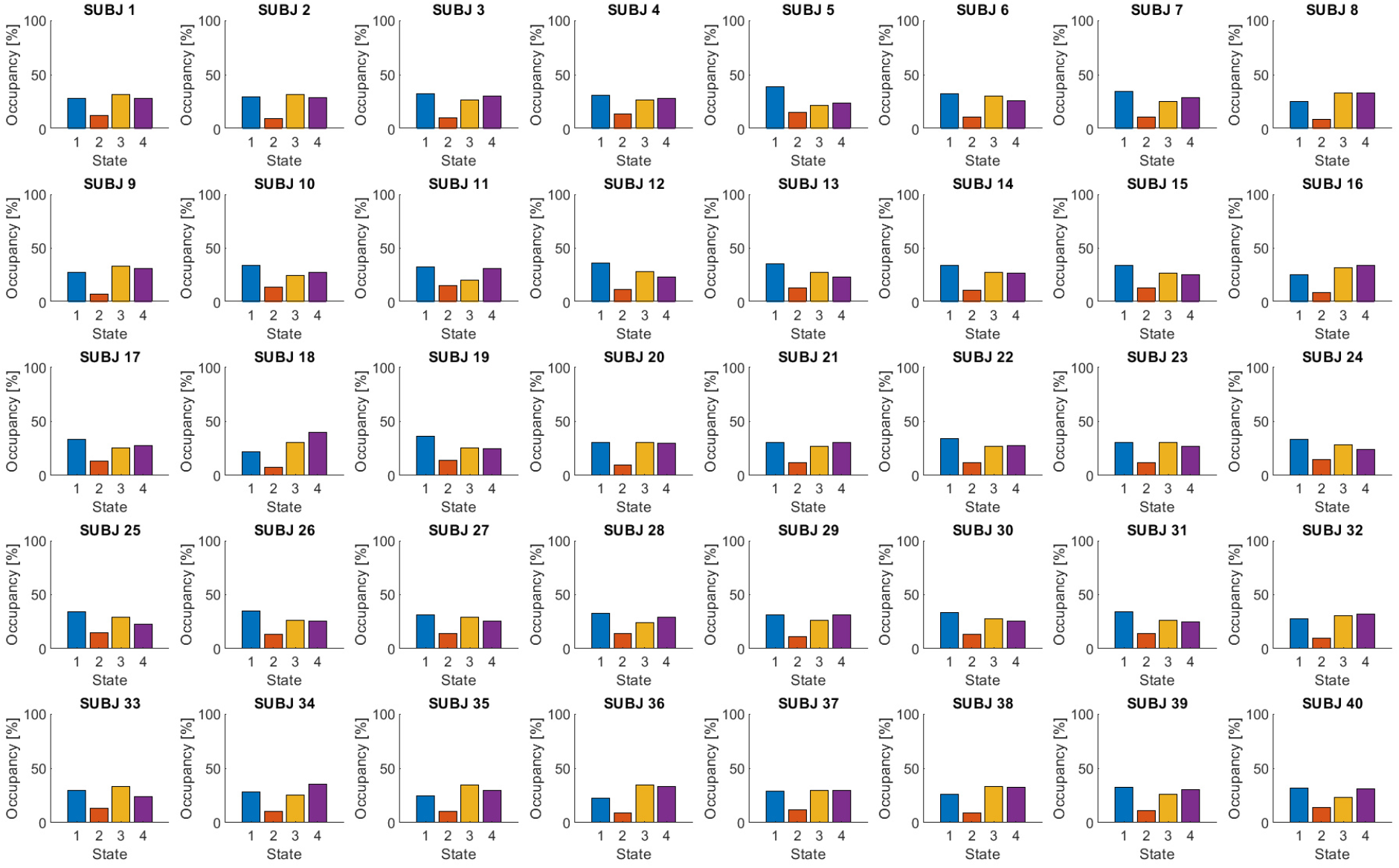
Fractional occupancy per subject in experiment 1. Each plot depicts the relative time spent in each of the four states of the final HMM for one of the subjects. The plots confirm that the states are evenly distributed across subjects, such that each subject spent some time in each of the states.

**Supplementary Figure 4:**
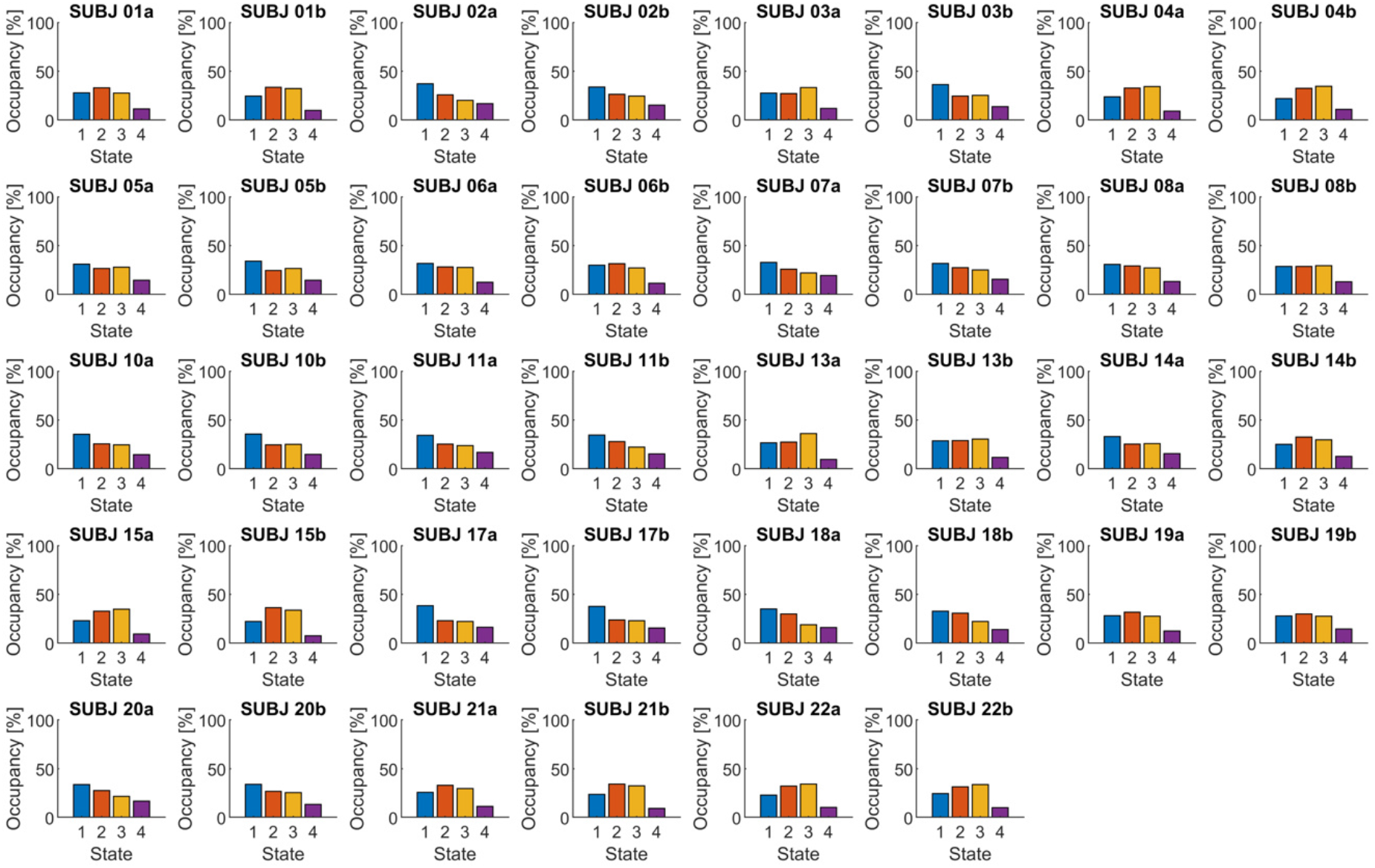
Fractional occupancy per subject in experiment 2. Each plot depicts the relative time spent in each of the four states of the final HMM for one of the experimental sessions of a subject. The plots confirm that the states are evenly distributed across subjects, such that each subject spent some time in each of the states.

### Supplementary Tables

**Supplementary Table 1.**
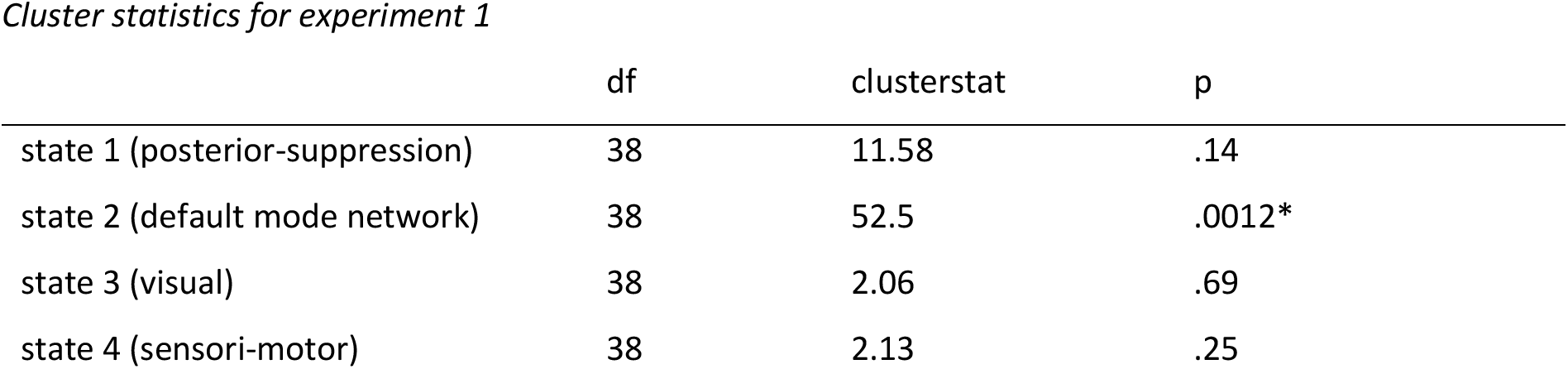
Cluster statistics for experiment 1. *Notes*. Results of independent samples random permutation cluster t-tests comparing the relative power increase to baseline between the tACS and the sham group separately for each state. Cluster test results received an additional Bonferroni correction for four multiple comparisons.

|  | df | clusterstat | p |
| --- | --- | --- | --- |
| state 1 (posterior-suppression) | 38 | 11.58 | .14 |
| state 2 (default mode network) | 38 | 52.5 | .0012* |
| state 3 (visual) | 38 | 2.06 | .69 |
| state 4 (sensori-motor) | 38 | 2.13 | .25 |

**Supplementary Table 2.** Repeated measures ANOVA in the theta and beta range in experiment 1. *Notes*. Repeated measures ANOVA in the individual theta (IAF-7 – IAF-3) and beta (IAF+4 – IAF+20) range. In contrast to results in the alpha band, neither an effect of stimulation nor a STIMULATION*STATE interaction is evident. However, there seems to be an overall difference in oscillatory power across states. Further there is a noteworthy trend for a STIMULATION*STATE interaction in the beta band, which could, however, not be replicated in the second experiment (Supplementary Table 4).

|  | df<sub>numerator</sub> | df<sub>denominator</sub> | F | p |
| --- | --- | --- | --- | --- |
| Theta (IAF-7 – IAF-3) |  |  |  |  |
| STIMULATION | 1 | 38 | 0.66 | .42 |
| STATE | 3 | 114 | 11.60 | <.001* |
| STIMULATION*STATE | 3 | 114 | 0.092 | .8 |
| Beta (IAF+4 – IAF+20) |  |  |  |  |
| STIMULATION | 1 | 38 | 1.23 | .27 |
| STATE | 3 | 114 | 13.18 | <.001* |
| STIMULATION*STATE | 3 | 114 | 2.28 | .08 |

**Supplementary Table 3.** Cluster statistics for experiment 2. *Notes*. Results of dependent samples random permutation cluster t-tests comparing the relative power increase to baseline between the tACS and the sham session separately for each state. Cluster test results received an additional Bonferroni correction for the four comparisons. Asterisks indicate significant differences.

| state | df | clusterstat | p |
| --- | --- | --- | --- |
| state 1 (posterior-suppression) | 18 | 4.72 | .46 |
| state 2 (visual) | 18 | 3.75 | .61 |
| state 3 (sensori-motor) | 18 | 14.82 | .066 |
| state 4 (default mode network) | 18 | 48.51 | .004* |

**Supplementary Table 4.** Repeated measures ANOVA in the theta and beta range in experiment 2. *Notes*. Repeated measures ANOVA in the individual theta (IAF-7 – IAF-3) and beta (IAF+4 – IAF+20) range. In contrast to results in the alpha band, neither an effect of stimulation nor a STIMULATION*STATE interaction is evident. However, there seems to be an overall difference in oscillatory power across states. The trend observed in the beta band, observed in experiment 1 is not replicated.

|  |  | df<sub>nominator</sub> | df<sub>denominator</sub> | F | p |
| --- | --- | --- | --- | --- | --- |
| Theta (IAF-7 – IAF-3) |  |  |  |  |  |
|  | STIMULATION | 1 | 18 | 2.96 | .1 |
|  | STATE | 3 | 54 | 12.36 | <.001* |
|  | STIMULATION*STATE | 3 | 54 | 1.53 | .23 |
| Beta (IAF+4 – IAF+20) |  |  |  |  |  |
|  | STIMULATION | 1 | 18 | 0.27 | .61 |
|  | STATE | 3 | 54 | 4.69 | .02* |
|  | STIMULATION*STATE | 3 | 54 | 1.42 | .25 |

## References

1. Vosskuhl, J., Strüber, D. & Herrmann, C. S. Non-invasive Brain Stimulation: A Paradigm Shift in Understanding Brain Oscillations. Front. Hum. Neurosci. 12, 211 (2018).

2. Kasten, F. H. & Herrmann, C. S. Discrete sampling in perception via neuronal oscillations— Evidence from rhythmic, non-invasive brain stimulation. European Journal of Neuroscience **n/a**, (2020).

3. Mellin, J. M. et al. Randomized trial of transcranial alternating current stimulation for treatment of auditory hallucinations in schizophrenia. Eur. psychiatr. 51, 25–33 (2018).

4. Alexander, M. L. et al. Double-blind, randomized pilot clinical trial targeting alpha oscillations with transcranial alternating current stimulation (tACS) for the treatment of major depressive disorder (MDD). Transl Psychiatry 9, 106 (2019).

5. Elyamany, O., Leicht, G., Herrmann, C. S. & Mulert, C. Transcranial alternating current stimulation (tACS): from basic mechanisms towards first applications in psychiatry. Eur Arch Psychiatry Clin Neurosci (2020) doi:10.1007/s00406-020-01209-9.

6. Fröhlich, F. & McCormick, D. A. Endogenous Electric Fields May Guide Neocortical Network Activity. Neuron 67, 129–143 (2010).

7. Krause, M. R., Vieira, P. G., Csorba, B. A., Pilly, P. K. & Pack, C. C. Transcranial alternating current stimulation entrains single-neuron activity in the primate brain. Proc Natl Acad Sci USA 116, 5747–5755 (2019).

8. Johnson, L. et al. Dose-dependent effects of transcranial alternating current stimulation on spike timing in awake nonhuman primates. Science Advances 6, eaaz2747 (2020).

9. Herrmann, C. S., Rach, S., Neuling, T. & Strüber, D. Transcranial alternating current stimulation: a review of the underlying mechanisms and modulation of cognitive processes. Front. Hum. Neurosci. 7, (2013).

10. Vossen, A., Gross, J. & Thut, G. Alpha Power Increase After Transcranial Alternating Current Stimulation at Alpha Frequency (α-tACS) Reflects Plastic Changes Rather Than Entrainment. Brain Stimulation 8, 499–508 (2015).

11. Zaehle, T., Rach, S. & Herrmann, C. S. Transcranial Alternating Current Stimulation Enhances Individual Alpha Activity in Human EEG. PLoS ONE 5, e13766 (2010).

12. Kasten, F. H., Dowsett, J. & Herrmann, C. S. Sustained Aftereffect of α-tACS Lasts Up to 70 min after Stimulation. Front. Hum. Neurosci. 10, (2016).

13. Neuling, T., Rach, S. & Herrmann, C. S. Orchestrating neuronal networks: sustained after-effects of transcranial alternating current stimulation depend upon brain states. Front. Hum. Neurosci. 7, (2013).

14. Wischnewski, M. et al. NMDA Receptor-Mediated Motor Cortex Plasticity After 20 Hz Transcranial Alternating Current Stimulation. Cerebral Cortex 29, 2924–2931 (2019).

15. Horvath, J. C., Forte, J. D. & Carter, O. Evidence that transcranial direct current stimulation (tDCS) generates little-to-no reliable neurophysiologic effect beyond MEP amplitude modulation in healthy human subjects: A systematic review. Neuropsychologia 66, 213–236 (2015).

16. Horvath, J. C., Forte, J. D. & Carter, O. Quantitative Review Finds No Evidence of Cognitive Effects in Healthy Populations From Single-session Transcranial Direct Current Stimulation (tDCS). Brain Stimulation 8, 535–550 (2015).

17. Vöröslakos, M. et al. Direct effects of transcranial electric stimulation on brain circuits in rats and humans. Nat Commun 9, 483 (2018).

18. Lafon, B. et al. Low frequency transcranial electrical stimulation does not entrain sleep rhythms measured by human intracranial recordings. Nat Commun 8, 1199 (2017).

19. Ridding, M. C. & Ziemann, U. Determinants of the induction of cortical plasticity by non-invasive brain stimulation in healthy subjects: Induction of cortical plasticity by non-invasive brain stimulation. The Journal of Physiology 588, 2291–2304 (2010).

20. Kasten, F. H., Duecker, K., Maack, M. C., Meiser, A. & Herrmann, C. S. Integrating electric field modeling and neuroimaging to explain inter-individual variability of tACS effects. Nat Commun 10, 5427 (2019).

21. Laakso, I., Tanaka, S., Koyama, S., De Santis, V. & Hirata, A. Inter-subject Variability in Electric Fields of Motor Cortical tDCS. Brain Stimulation 8, 906–913 (2015).

22. Riddle, J. et al. Brain-derived neurotrophic factor (BDNF) polymorphism may influence the efficacy of tACS to modulate neural oscillations. Brain Stimulation 13, 998–999 (2020).

23. Ruhnau, P. et al. Eyes wide shut: Transcranial alternating current stimulation drives alpha rhythm in a state dependent manner. Sci Rep 6, 27138 (2016).

24. Feurra, M. et al. State-Dependent Effects of Transcranial Oscillatory Currents on the Motor System: What You Think Matters. Journal of Neuroscience 33, 17483–17489 (2013).

25. Alagapan, S. et al. Modulation of Cortical Oscillations by Low-Frequency Direct Cortical Stimulation Is State-Dependent. PLoS Biol 14, e1002424 (2016).

26. Feurra, M. et al. State-Dependent Effects of Transcranial Oscillatory Currents on the Motor System during Action Observation. Scientific Reports 9, 12858 (2019).

27. Vidaurre, D. et al. Spontaneous cortical activity transiently organises into frequency specific phase-coupling networks. Nat Commun 9, 2987 (2018).

28. Baker, A. P. et al. Fast transient networks in spontaneous human brain activity. eLife 3, e01867 (2014).

29. Vidaurre, D. et al. Spectrally resolved fast transient brain states in electrophysiological data. NeuroImage 126, 81–95 (2016).

30. Quinn, A. J. et al. Unpacking Transient Event Dynamics in Electrophysiological Power Spectra. Brain Topogr 32, 1020–1034 (2019).

31. Kasten, F. H. & Herrmann, C. S. Recovering Brain Dynamics During Concurrent tACS-M/EEG: An Overview of Analysis Approaches and Their Methodological and Interpretational Pitfalls. Brain Topogr 32, 1013–1019 (2019).

32. Veniero, D., Vossen, A., Gross, J. & Thut, G. Lasting EEG/MEG Aftereffects of Rhythmic Transcranial Brain Stimulation: Level of Control Over Oscillatory Network Activity. Front. Cell. Neurosci. 9, (2015).

33. Neuling, T., Wagner, S., Wolters, C. H., Zaehle, T. & Herrmann, C. S. Finite-Element Model Predicts Current Density Distribution for Clinical Applications of tDCS and tACS. Front. Psychiatry 3, (2012).

34. Kasten, F. H. & Herrmann, C. S. Transcranial Alternating Current Stimulation (tACS) Enhances Mental Rotation Performance during and after Stimulation. Front. Hum. Neurosci. 11, (2017).

35. Oldfield, R. C. The assessment and analysis of handedness: The Edinburgh inventory. Neuropsychologia 9, 97–113 (1971).

36. Oostenveld, R., Fries, P., Maris, E. & Schoffelen, J.-M. FieldTrip: Open Source Software for Advanced Analysis of MEG, EEG, and Invasive Electrophysiological Data. Computational Intelligence and Neuroscience 2011, 1–9 (2011).

37. Taulu, S. & Simola, J. Spatiotemporal signal space separation method for rejecting nearby interference in MEG measurements. Phys. Med. Biol. 51, 1759–1768 (2006).

38. Nenonen, J. et al. Validation of head movement correction and spatiotemporal signal space separation in magnetoencephalography. Clinical Neurophysiology 123, 2180–2191 (2012).

39. Taulu, S., Simola, J. & Kajola, M. Applications of the signal space separation method. IEEE Trans. Signal Process. 53, 3359–3372 (2005).

40. Van Veen, B. D., Van Drongelen, W., Yuchtman, M. & Suzuki, A. Localization of brain electrical activity via linearly constrained minimum variance spatial filtering. IEEE Trans. Biomed. Eng. 44, 867–880 (1997).

41. Colclough, G. L., Brookes, M. J., Smith, S. M. & Woolrich, M. W. A symmetric multivariate leakage correction for MEG connectomes. NeuroImage 117, 439–448 (2015).

42. Baum, L. E., Petrie, T., Soules, G. & Weiss, N. A Maximization Technique Occurring in the Statistical Analysis of Probabilistic Functions of Markov Chains. Ann. Math. Statist. 41, 164–171 (1970).

43. Vidaurre, D. et al. Discovering dynamic brain networks from big data in rest and task. NeuroImage 180, 646–656 (2018).

44. Pohle, J., Langrock, R., van Beest, F. M. & Schmidt, N. M. Selecting the Number of States in Hidden Markov Models: Pragmatic Solutions Illustrated Using Animal Movement. JABES 22, 270–293 (2017).

